# Biomimetic Cascade-Targeting Drug Delivery System for Reversing Chemoresistance in Osteosarcoma

**DOI:** 10.1101/2025.11.04.686585

**Authors:** Xingyin Li, Shaojie Xu, Youyun Peng, Ziliang Su, Sining Wang, Ang Li, Yaying Du, Zengwu Shao, Xin Huang

**Author notes:** Corresponding author: Xin Huang, Zengwu Shao and Yaying Du. Xingyin Li, Shaojie Xu and Youyun Peng contributed equally.

## Abstract

Osteosarcoma (OS) remains the most prevalent primary malignant bone tumor in adolescents, with chemotherapy resistance significantly limiting the efficacy of current treatments. To address the challenges of chemoresistance driven by DNA repair mechanisms and drug efflux, we developed a biomimetic cascade-targeting drug delivery system, TAT-mPDO@cRGD-M. This system combines a polydopamine (mPDA) core with strong photothermal and drug-loading capacities, enabling the co-delivery of cisplatin (CDDP) and olaparib (OLA). Surface functionalization with the nuclear localization peptide TAT and cloaking with osteosarcoma cell membranes modified with cRGD peptides allows for selective tumor recognition, nuclear targeting, and stimuli-responsive drug release. Upon near-infrared (NIR) irradiation, the platform enhances DNA damage, suppresses the PI3K-AKT signaling pathway, and promotes apoptosis while inhibiting epithelial-mesenchymal transition (EMT). In vitro and in vivo studies demonstrated that TAT-mPDO@cRGD-M exhibits potent antitumor activity against cisplatin-resistant osteosarcoma, maintaining strong therapeutic efficacy under low-power NIR irradiation without causing significant damage to normal tissues. This study highlights a highly specific and biocompatible therapeutic approach that offers promising potential for overcoming chemotherapy resistance in osteosarcoma and advancing clinical treatment strategies.

## Introduction

Osteosarcoma (OS) is the most prevalent primary malignant bone tumor in children and adolescents, with an annual incidence of approximately 3 cases per million^1^. OS shows high aggressiveness, with about 20% of patients presenting with pulmonary metastases at initial diagnosis and 35-45% experiencing recurrence and poor prognosis^2,3^. Surgical resection alone yields a 5-year survival rate of merely 15-20%^4^. Chemotherapy remains the principal adjuvant strategy, among which CDDP is one of the most widely used agents. Combined surgical and chemotherapeutic regimens have improved the 5-year survival rate to around 75%^5^. However, over the past three and a half decades, no breakthrough progress has been achieved in OS chemotherapy^6^ and the long-term survival rate for patients with metastatic or recurrent disease remains below 20%^7^.

Chemoresistance represents one of the major obstacles limiting the improvement of survival outcomes in OS patients. The therapeutic efficacy of CDDP is markedly compromised by intrinsic or acquired resistance, and CDDP exhibits only limited efficacy when administered as a single agent^8^. Such resistance may be attributable to reduced intracellular accumulation, enhanced efflux, or upregulated DNA repair capacity^9^. Moreover, osteosarcoma often exhibits significant chromosomal instability^10,11^, with most tumors carrying mutations resembling those seen in BRCA1/2-deficient malignancies^12^. These features suggest that osteosarcoma may rely on certain DNA damage repair pathways, providing a theoretical basis for utilizing synthetic lethality strategies.

The combination of the poly ADP-ribose polymerase (PARP) inhibitor OLA with DNA-damaging chemotherapy has been considered a promising anti-osteosarcoma strategy. Park et al. demonstrated in murine and cellular models that olaparib synergized with doxorubicin to inhibit OS growth and induce apoptosis^13^. Liang et al. reported that olaparib-mediated PARP1 inhibition restored cisplatin sensitivity in osteosarcoma cells and organoids^14^. However, available clinical data suggest that cisplatin combined with olaparib yields suboptimal efficacy^15,16^, potentially due to insufficient accumulation of drugs within the nucleus where they exert their pharmacological action. Only a small fraction of intracellular cisplatin molecules can passively diffuse into the nucleus^17^; cisplatin molecules that fail to reach the nucleus are readily pumped out of osteosarcoma cells by ATP-binding cassette (ABC) family efflux transporters, which are often overexpressed on the OS cell membrane^18^. The trans-activator of transcription (TAT) peptide has been demonstrated to be an effective nuclear-targeting agent, enabling spatiotemporal proximity to DNA^19^. A major limitation for TAT application is its non-specific cellular internalization^20^.

Engineered membrane-coated nanoparticles have been extensively studied for tumor-targeted delivery^21^. The cell membrane coating exerts a spatial steric shielding effect on TAT peptides, thereby reducing the risk of premature endocytosis or aberrant distribution. In our preliminary studies^22^, cRGD-modified tumor cell membrane-coated nanoparticles selectively targeted and were taken up by osteosarcoma cells by targeting the αvβ3 integrin, which is highly expressed in osteosarcoma^23,24^. Additionally, the cRGD peptide enhances vascular and tissue permeability by targeting ganglioside-1, facilitating deeper tissue penetration and improved therapeutic delivery^25^. Furthermore, membrane-coated nanodrugs are taken up via membrane fusion, thus bypassing endocytic/lysosomal degradation pathways and enabling direct cytoplasmic delivery^26^. This mechanism effectively circumvents lysosomal destruction of TAT-modified nanoparticle cores before they can exert its nuclear localization function. Therefore, synergistic delivery of therapeutics via cRGD-modified membrane coating and TAT-modified nanoparticle cores represents a promising cascade-targeting strategy for osteosarcoma treatment.

In this study, a cascade-targeting nanodelivery system named TAT-mPDO@cRGD-M was constructed. After specific recognition and uptake by tumor cells, the nanodelivery system facilitated the nuclear localization of mPDO under the guidance of TAT peptides (Fig. 1). This study is the first to evaluate the correlation between PARP1 and PARP2 expression patterns and chemotherapy resistance in osteosarcoma using clinical samples. By reversing cisplatin resistance in osteosarcoma cells with olaparib, it provides new insights for the clinical treatment of chemotherapy-resistant osteosarcoma patients.

**Fig. 1.**
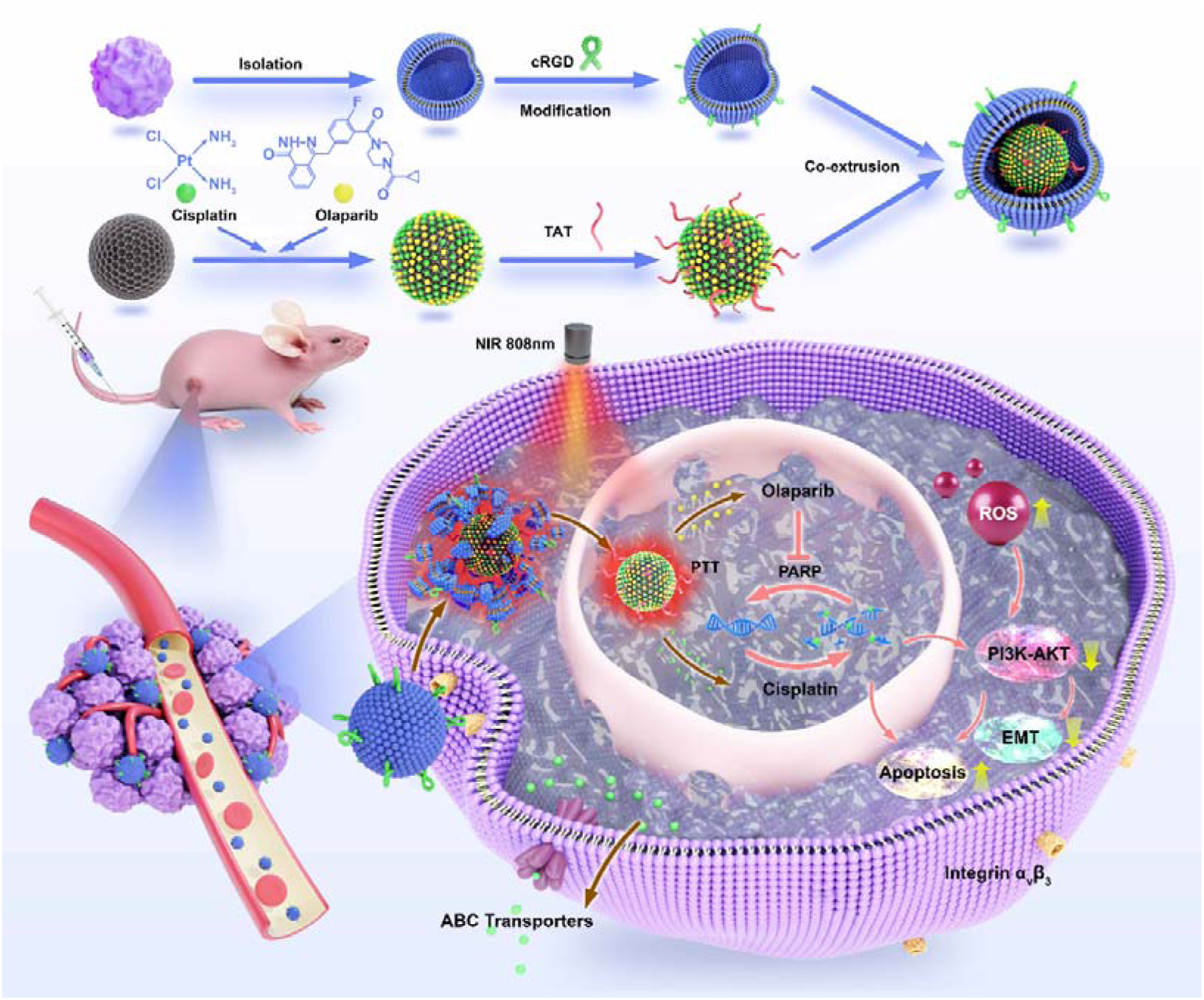
The synthesis, nuclear delivery, and antitumor mechanism of TAT-mPDO@cRGD-M. Cisplatin and olaparib were co-loaded into mPDO, followed by surface modification with TAT and coating with a cRGD-modified tumor cell membrane. The biomimetic nanoplatform enables tumor targeting and nuclear co-delivery of drugs, leading to PARP inhibition, enhanced DNA damage, ROS generation, and suppression of the PI3K/AKT pathway, thereby inhibiting EMT, promoting apoptosis, and achieving effective tumor suppression.

## Materials and Methods

### Establishment of stable cisplatin-resistant osteosarcoma cell lines

MNNG/HOS C1#5 [R-1059-D] (CL-0492) and U2OS (CL-0236) were kindly provided by Procell (Hubei, China), RRIDs, CVCL_0439 and CVCL_0042 respectively) and cultured in MEM and McCoy’s 5A medium (Procell, Hubei, China), respectively, supplemented with 10% FBS (Thermo Fisher, Waltham, MA, USA). MNNG/HOS and U2OS cells were exposed to a gradient of cisplatin concentrations for 48 h. Cell viability was assessed using the CCK-8 assay. After incubation with the working solution in the dark, absorbance was measured, and the IC_50_ value of cisplatin was calculated. Subsequently, cells were cultured in medium containing cisplatin at their respective IC_50_ concentrations. After five passages, the cisplatin concentration was doubled, and the cells were continuously passaged five more times. This stepwise selection process was repeated with gradually increased cisplatin concentrations until stable MNNG/HOS and U2OS cell lines were obtained that could proliferate steadily in medium containing five times their initial IC_50_ concentration of cisplatin.

### Synthesis and characterization of the nanodelivery system

mPDA (Ruixitech, Cat# R-MPDA) and cRGD (Ruixitech, Cat# R-9998) was purchased from Xi’an Ruixi Biologicals, while CDDP (MedChemExpress, Cat# HY-17394), OLA (MedChemExpress, Cat# HY-10162) and TAT (MedChemExpress, Cat# HY-P0281) were from MedChemExpress. Equal amounts of CDDP and OLA were dissolved in water and ethanol, respectively, and then mixed with the same amount of mPDA solution. The mixture was stirred overnight in an ice bath, centrifuged at 8000 rpm for 10 min, and the precipitate was collected, washed with water, and redispersed to obtain mPDO. Using the same procedure, mPD and TAT-mPDO were prepared. Tumor cell membranes were isolated from MNNG/HOS-CDDP and U2OS-CDDP cells using the Minute™ Plasma Membrane Kit (MedChemExpress, Cat# HY-17394). Extracted tumor cell membranes were mixed with cRGD solution at the optimal ratio and incubated at 40°C for 2 hours, followed by repeated washing to obtain the cRGD-modified tumor cell membranes. The membranes were sequentially extruded through 400nm and 200nm polycarbonate membranes for approximately 20 cycles to obtain cRGD-M. mPDO@cRGD-M and TAT-mPDO@cRGD-M were prepared by co-extruding the nanoparticles with cRGD-modified tumor cell membranes. mPDO, cRGD-M and TAT-mPDO@cRGD-M solutions were sonicated for 5 minutes, separately applied to copper grids, and negatively stained with uranyl acetate for observation using a low-voltage transmission electron microscope (TEM) system (Hitachi, Japan).

The particle size distribution and zeta potential of mPDA, mPDO, TAT-mPDO, cRGD-M and TAT-mPDO@cRGD-M were analyzed by dynamic light scattering (DLS, Malvern-Zetasizer, UK). The particle sizes of mPDO, TAT-mPDO and TAT-mPDO@cRGD-M in physiological saline were monitored over 48 hours. After mixed with fetal bovine serum (FBS), absorbance at 540 nm was measured at 3, 6, 12, 24 and 48 hours to assess serum stability. Additionally, hemolysis was evaluated by incubating mouse red blood cell suspensions with PBS (negative control), deionized water (positive control), or serially diluted TAT-mPDO@cRGD-M solutions for 3 hours, followed by centrifugation and measurement of supernatant absorbance at 540 nm. Hemolysis percentage was normalized using PBS as 0% and H2O as 100% according to the formula:

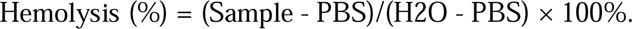

MNNG/HOS-CDDP cells membrane, cRGD-M and nanoparticles were analyzed by sodium dodecyl sulfate-polyacrylamide gel electrophoresis (SDS-PAGE). Briefly, protein samples were prepared by mixing with loading buffer, then heated at 100°C for 10min. The samples were loaded onto a polyacrylamide gel and subjected to electrophoresis. The protein bands were then stained with Coomassie Blue.

Solutions of cisplatin and olaparib were prepared with defined concentration gradients, and the peak areas of cisplatin and olaparib at different concentrations were determined by high-performance liquid chromatography (HPLC) to generate standard curves. During mPDO synthesis, the supernatant and wash solutions were collected after three washes, and drug content was quantified using the corresponding standard curves. The drug encapsulation efficiency and drug loading efficiency were calculated as follow:

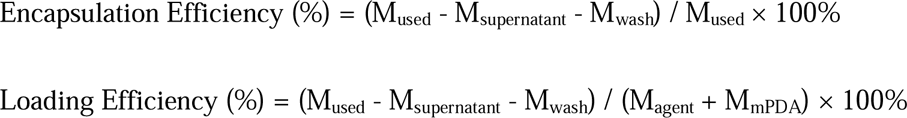

where M_used_ is the total amount (mg) of drug initially added, M_supernatant_ and M_wash_ are the drug amounts (mg) in the supernatant and wash solution, M_agent_ is the weight(mg) of CDDP or OLA encapsulated inside the nanoparticles and M_mPDA_ is the total amount (mg) of mPDA used in nanoparticle synthesis.

The drug release efficiency was assessed under conditions with or without NIR irradiation and low pH, using the same method. For acidic conditions, drug-loaded samples were incubated in PBS at pH 5.0 or pH 7.4. Cumulative release percentage (%) was defined using the following formulas:

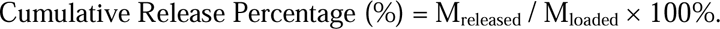

where M_released_ represents the total weight (mg) of the released drug and M_loaded_ is the total weight (mg) of the initial loaded drug.

For evaluating the photothermal performance of the material, both in vitro and in vivo methods were employed. In vitro, TAT-mPDO@cRGD-M solution was irradiated with an 808 nm near-infrared laser at 2 W/cm² for 2 minutes, followed by a 2-minute pause, with temperature changes recorded using an infrared thermal imager (Testo 865; Testo, Schwarzwald, Germany). In the tumor-bearing nude mouse model, TAT-mPDO@cRGD-M was injected via the tail vein, followed by three 2-minute sessions of 808 nm NIR irradiation at 2 W/cm² on next day, tumor temperature was recorded throughout. The in vivo photothermal effect was also assessed with the NIR irradiation at 1 W/cm² using the same procedure.

### Cell functional assays

MNNG/HOS-CDDP cells were treated with 10 μg/mL of PBS, mPD, mPDO, TAT-mPDO, mPDO@cRGD-M, or TAT-mPDO@cRGD-M for 12, 24, or 48 hours, or with 5, 10, or 20 μg/mL of above materials for 48 hours. Simultaneously, 808 nm NIR irradiation were applied at 2 W/cm². Cell viability was assessed using the CCK-8 assay (MedChemExpress, Cat# HY-K0301). The live/dead and migration assays were performed using the same treatment groups (10μg/mL, 48h) as the CCK-8 assay. For live/dead staining, cells were stained using a Calcein-AM/propidium iodide kit (Beyotime, Cat# C1371S). For the migration assay, pretreated cells were resuspended in 100 μL serum-free medium and seeded into the upper chambers of Transwell inserts (lower chambers, 10% FBS). After 12-16 hours, migrated cells were fixed and stained. The same procedures were applied to U2OS-CDDP cells.

### Flow Cytometry Analysis

MNNG/HOS-CDDP cells were treated with TAT-mPDO@cRGD-M (0, 2, 5, 10, and 20 μg/mL for 48 h) for ROS measurement, and PBS, mPD, or mPDO (10 μg/mL for 48 h) for apoptosis detection, all under 808 nm NIR irradiation at 2 W/cm². ROS levels were measured using the DCFH-DA Assay Kit (Beyotime, Cat# S0035S) followed by flow cytometry. Apoptosis was detected using the Annexin V-FITC/PI Apoptosis Detection Kit (Vazyme, Cat# A211).

For ROS measurement, cells were harvested, washed with cold PBS, and incubated with 10 μM DCFH-DA at 37°C for 30 minutes. After washing with PBS, cells were resuspended in PBS and analyzed by flow cytometry. Fluorescence signals were collected, and data were analyzed using FlowJo software to quantify fluorescence intensity. For apoptosis detection, cells were harvested, washed with cold PBS, and stained with 5 μL of Annexin V-FITC and 5 μL of PI at room temperature for 10 minutes, protected from light. After adding 400 μL of Binding Buffer, cells were analyzed by flow cytometry within 1 hour. A minimum of 10,000 events were recorded per sample, and appropriate controls were used for compensation and gating. FlowJo software was used for data analysis and quantification of fluorescence intensity.

### Immunofluorescence staining

mPDO and TAT-mPDO were labeled with green fluorescent FITC and co-extruded with cRGD-M to obtain FITC-labeled mPDO@cRGD-M and TAT-mPDO@cRGD-M. MNNG/HOS-CDDP cells (5000 cells/well) were seeded in confocal dish plates, and after cell attachment, serum-free medium was added, followed by the addition of 10 μL of mPDO@cRGD-M and TAT-mPDO@cRGD-M at the same concentration. The cells were incubated at 37°C for 4 hours, and images were acquired using confocal laser scanning microscopy (CLSM).

FITC-labeled mPDO, TAT-mPDO, mPDO@cRGD-M and TAT-mPDO@cRGD-M were intravenously injected via the tail vein into tumor-bearing nude mice. On next day, the biodistribution of nanoparticles was assessed using an in vivo imaging system. Then mice were euthanized, and their vital organs (heart, liver, spleen, lungs, kidneys) and tumors were extracted for fluorescence imaging.

### Animal Models

BALB/c Nude mice (male, 3 weeks) were obtained from GemPharmatech (Guangdonog, China) and maintained under SPF conditions. All procedures were approved by the Animal Ethics Committee of Union Hospital, Tongji Medical College (Huazhong University of Science and Technology, Wuhan, China).

Male BALB/c nude mice were subcutaneously injected with MNNG/HOS-CDDP tumor cells (3-4 × 10^4^ cells/μL). When the tumor diameter reached 5-6 mm, the mice were randomly divided into six groups: control, mPD, mPDO, TAT-mPDO, mPDO@cRGD-M, and TAT-mPDO@cRGD-M. Each group was injected via the tail vein with different drugs at the same concentration. 24 h after injection, the tumor site was irradiated with an 808 nm NIR laser (2 W/cm^2^) for 2 minutes, repeated three times. Tumor volume and body weight were monitored daily. On day 18, all mice were euthanized, and blood was collected via the orbital venous. Tumors, heart, liver, spleen, lungs, and kidneys were also harvested. Blood biochemical parameters were assessed. After photographing and weighing the tumors, the organs and tumors were embedded in paraffin and sectioned. Organs were stained with hematoxylin and eosin (H&E), and tumors were stained with H&E and IHC tested for Ki67 (Ki67 Rabbit mAb, Abclonal, Cat# A20018), CD31 (CD31/PECAM1 Rabbit mAb, Abclonal, Cat# A19014), Cleaved Caspase-3 (active Caspase-3 Rabbit mAb, Abclonal, Cat# A11021), Cleaved PARP1 (Cleaved PARP1 p25 Rabbit mAb, Abclonal, Cat# A19612), p-AKT (p-AKT-S473 Rabbit mAb, Abclonal, Cat# AP0637), and p-H2AX (Phospho-Histone H2AX-S139 Rabbit mAb, Abclonal, Cat# AP1555).

Additionally, MNNG/HOS-CDDP-loaded tumor-bearing mice were randomly divided into four groups: mPDO, mPDO+Laser, TAT-mPDO@cRGD-M, and TAT-mPDO@cRGD-M+Laser. In contrast to the previous method, NIR irradiation with a power of 1 W/cm^2^ was applied to the tumor site the day after drug injection, followed by another drug injection every 3 days. Tumor size was monitored daily. On day 15, the experiment was terminated, and mice were euthanized. Tumor tissues were collected for photographing and weighing.

### Immunohistochemistry (IHC) analysis

Tumor tissue sections were incubated with appropriate primary antibodies at 4°C for 20 h, followed by incubation with secondary antibody for 1 h. After visualization using diaminobenzidine (DAB) solution and counterstaining with hematoxylin, images were captured under a microscope and analyzed using ImageJ software.

### RNA Sequencing

MNNG/HOS and MNNG/HOS-CDDP cells, as well as MNNG/HOS-CDDP cells treated with PBS and 10 μg/mL TAT-mPDO@cRGD-M for 48 hours, were collected for transcriptome sequencing on the Illumina platform. Total RNA was extracted, cDNA libraries were constructed and purified, and RNA-seq was performed for in-depth analysis. Three biological replicates per group were included. The alignment rate of each sample to the reference genome exceeded 95%, ensuring high-quality sequencing data (Table S1, S2).

### Reverse transcriptase quantitative polymerase chain reaction (RT-qPCR)

MNNG/HOS-CDDP cells treated with PBS and 10 μg/mL TAT-mPDO@cRGD-M for 48 hours were collected. Total RNA was extracted using the RNeasy Mini Kit, and cDNA was synthesized using the iScript cDNA Synthesis Kit. RT-qPCR was performed using SYBR Premix Ex Taq and the MxPro Mx3005P real-time PCR detection system (Agilent Technologies, Santa Clara, CA). Primer sequences are listed in Table S3. GAPDH was used as an internal control.

### Western blot

MNNG/HOS-CDDP and U2OS-CDDP cells were treated with TAT-mPDO@cRGD-M (10 μg/mL) for 48 hours. Additionally, MNNG/HOS-CDDP cells were first treated with 20 μM 740Y-P (PI3K activator, MedChemExpress, Cat# HY-P0175) for 1 hour, followed by the addition of TAT-mPDO@cRGD-M under the same conditions. Simultaneously, MNNG/HOS cells were treated with 10 μg/mL mPDO and TAT-mPDO for 48 hours. The drug-treated cells were collected and lysed on ice for 20 minutes using 1× RIPA buffer supplemented with 1% PMSF. Protein concentration was determined using a BCA assay kit (Boster, Hubei, China). Protein supernatant was mixed with 5× loading buffer at a 4:1 volume ratio and heated at 95°C for 10 minutes. 20 μg of protein per lane was separated on SDS-PAGE and transferred to PVDF membranes. The membrane was blocked at room temperature with a rapid blocking buffer for 15-30 minutes, then incubated overnight at 4°C with appropriately diluted primary antibodies. After washing with TBST, the membrane was incubated with species-specific secondary antibodies for 30-60 minutes. Finally, Immunoreactive bands were visualized using an enhanced chemiluminescence (ECL) detection kit. The primary antibodies included MRP1/2, MDR1, p-H2AX, Cleaved Caspase3, Cleaved PARP1, BCL2, BAX, p-PI3K, PI3K, AKT, p-AKT, NCAD, MMP3, Vimentin and GAPDH (Abclonal, Thermo Fisher and Affinity). Goat anti-mouse/anti-rabbit secondary antibodies were purchased from Proteintech.

### Statistical Analyses

The data were analyzed using t tests, one-way ANOVA or Kruskal-Wallis test in Prism 9 (GraphPad, La Jolla, CA, USA). p < 0.05 was considered to indicate significance.

## Results and Discussion

### Clinical Significance of PARP in Chemoresistant Osteosarcoma

We analyzed tumor samples from 32 osteosarcoma patients to assess the clinical relevance of PARP expression. The median immunohistochemical (IHC) scores for PARP1 and PARP2 (3.2 and 6.1, respectively) were used as cutoff values for defining positive expression (Fig. S1A). Elevated PARP1 expression was significantly associated with advanced tumor stage (P = 0.010), chemoresistance (P = 0.012), and PARP2 positivity (P = 0.005) (Table 1). In contrast, PARP2 expression did not predict chemotherapy response. Receiver operating characteristic (ROC) curve analysis showed that PARP1 had an area under the curve (AUC) of approximately 0.70 for predicting chemotherapeutic responsiveness, indicating moderate discriminatory power (Fig. S1B).

**Table 1.** Clinicopathologic variables and the expression of PARP1/2 in 32 osteosarcomas.

| Characteristics |  | No. | PARP1<br>positive | P | PARP2<br>positive | P |
| --- | --- | --- | --- | --- | --- | --- |
| Age | ≤16 | 17 | 8(47%) | 0.723 | 7(41%) | 0.288 |
|  | > 16 | 15 | 8(53%) |  | 9(60%) |  |
| Sex | male | 21 | 10(48%) | 0.710 | 11(52%) | 0.710 |
|  | female | 11 | 6(55%) |  | 5(46%) |  |
| Tumor size,cm | ≤8 | 20 | 10(50%) | 1.000 | 9(45%) | 0.465 |
|  | >8 | 12 | 6(50%) |  | 7(58%) |  |
| Stage | Stage I | 2 | 1(50%) | 0.010 | 1(50%) | 0.432 |
|  | Stage II | 23 | 8(35%) |  | 10(44%) |  |
|  | Stage III-IV | 7 | 7(100%) |  | 5(71%) |  |
| Metastasis or<br>Recurrence | presence | 15 | 10(67%) | 0.077 | 8(53%) | 0.723 |
|  | absence | 17 | 6(35%) |  | 8(47%) |  |
| Histological grade | high | 30 | 15(50%) | 1.000 | 15(50%) | 1.000 |
|  | low | 2 | 1(50%) |  | 1(50%) |  |
| Histologic type | conventional | 29 | 15(52%) | 0.544 | 15(52%) | 0.544 |
|  | non-conventional | 3 | 1(33%) |  | 1(33%) |  |
| Chemosensitivity | Chemoresistant | 19 | 13(68%) | 0.012 | 11(58%) | 0.280 |
|  | chemosensitive | 13 | 3(23%) |  | 5(39%) |  |
| PARP2 | ≤6.1224 | 16 | 4(25%) | 0.005 |  |  |
|  | > 6.1224 | 16 | 15(75%) |  |  |  |

### Development of Resistant Cell Models

Although BRCA1/2 inactivation has been reported in 91% and 78% of OS tumors, respectively^27^, authentic BRCA1/2-deficient osteosarcoma cell lines remain rare^12^. MNNG/HOS cells harbor a PTEN loss-of-function mutation accompanied by ATM deficiency, a molecular phenotype considered to exhibit “BRCA-like” characteristics^12^. Both MNNG/HOS and U2OS cell lines, similar to primary OS tissues, express high levels of integrin α_v_β_3_^28,29^, making them ideal models for evaluating cRGD-modified membrane-coated nanodrugs. To establish cisplatin-resistant models, MNNG/HOS and U2OS cells were subjected to stepwise cisplatin selection, generating MNNG/HOS-CDDP and U2OS-CDDP derivatives. In U2OS cells, the IC₅₀ of cisplatin increased from 1.120 μg/mL in parental cells to 6.097 μg/mL in U2OS-CDDP cells, corresponding to a drug resistance index (DRI) of 5.43. Similarly, the IC₅₀ values in MNNG/HOS cells increased from 2.224 μg/mL to 9.671 μg/mL following resistance induction, yielding a DRI of 4.35 (Fig. S1C).

Transcriptomic sequencing was performed to compare MNNG/HOS and MNNG/HOS-CDDP cells, which identified 3,309 differentially expressed genes (DEGs), including 1,438 upregulated and 1,871 downregulated genes (Fig. S1D). Hierarchical clustering analysis demonstrated clear segregation between parental and resistant cells (Fig. S1E), confirming the transcriptional reprogramming associated with the acquisition of resistance. We further investigated genes previously reported to be associated with osteosarcoma chemoresistance^30^ and observed significant differential expression of these in MNNG/HOS-CDDP cells (Fig. S1F), thus supporting the successful establishment of a stable cisplatin-resistant model.

Previous studies have shown that PARP1 and PARP2 expression levels were elevated in certain cisplatin-resistant OS cell lines compared with their parental counterparts^31^; this upregulation was similarly observed in our resistant models (Fig. S1F). Gene set enrichment analysis (GSEA) further revealed significant enrichment of DNA damage response and DNA repair-related pathways in MNNG/HOS-CDDP cells (Fig. S1G), suggesting that enhanced DNA repair capacity may underlie the acquired cisplatin resistance phenotype.

### Fabrication and Characterization of TAT-mPDO@cRGD-M

The TAT-mPDO@cRGD-M nanoplatform was constructed through a three-step process: (1) co-loading of CDDP and OLA into mesoporous mPDA, followed by conjugation with TAT to obtain TAT-mPDO; (2) extraction of tumor cell membranes from MNNG/HOS-CDDP cells and functionalization with cRGD to generate cRGD-M; and (3) membrane coating of cRGD-M onto TAT-mPDO via serial extrusion to yield the final TAT-mPDO@cRGD-M.

Transmission electron microscopy (TEM) revealed that cRGD-M exhibited a characteristic vesicular morphology, whereas TAT-mPDO@cRGD-M displayed a distinct core-shell structure with an average diameter of approximately 100 nm, in which the cell membrane thickness was ∼10 nm (Fig. 2A). SDS-PAGE analysis demonstrated that the protein profiles of cRGD-M and TAT-mPDO@cRGD-M were highly consistent with those of native tumor cell membranes (Fig. 2G), indicating effective preservation of membrane-associated proteins, thus supporting the structural basis for homotypic targeting.

**Fig. 2.**
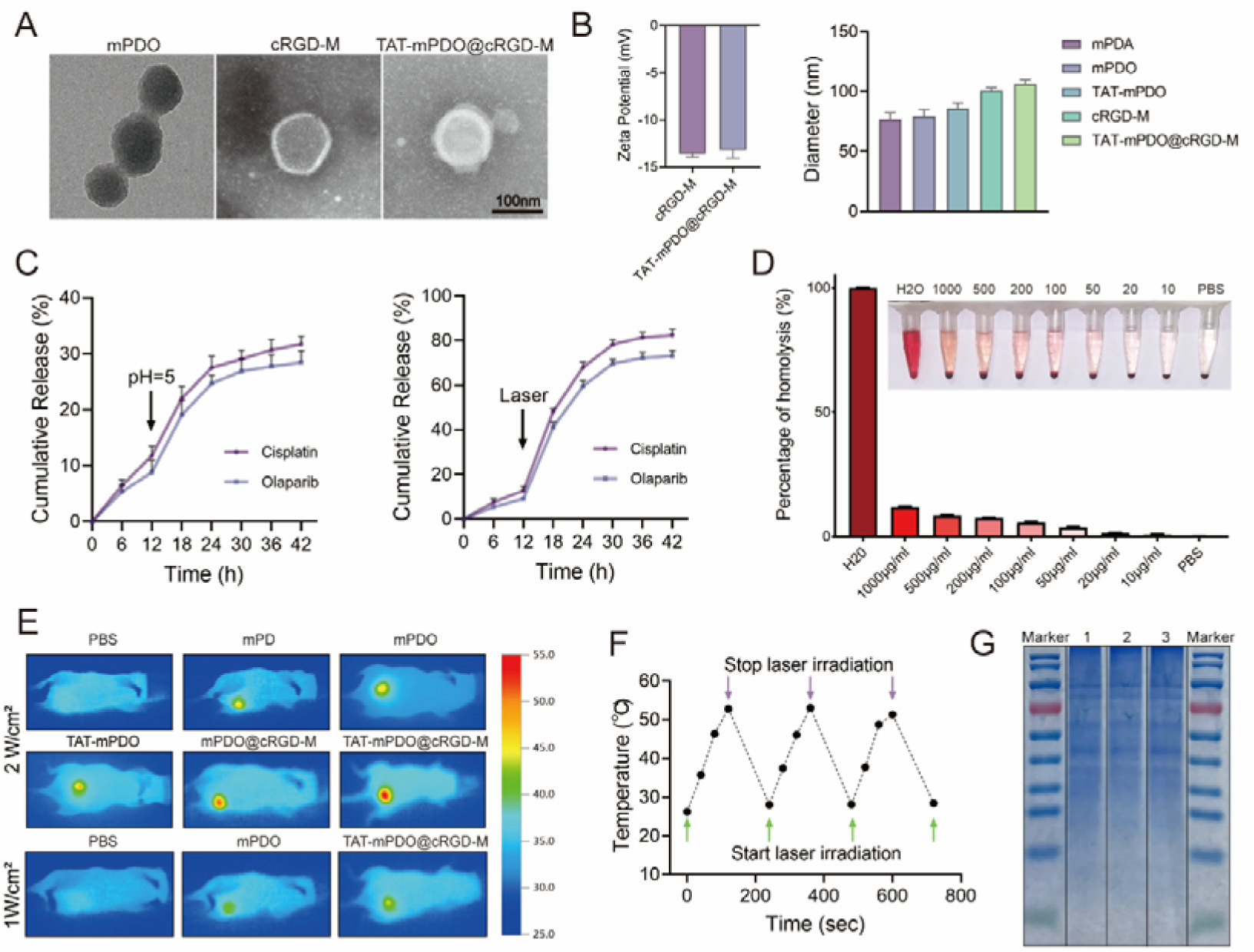
Characterizations of TAT-mPDO@cRGD-M. (A) TEM images showing the morphology of mPDO, cRGD-M, and TAT-mPDO@cRGD-M (scale bar: 100 nm). (B) Zeta potential of cRGD-M, TAT-mPDO@cRGD-M and particle size of mPDA, mPDO, TAT-mPDO, cRGD-M and TAT-mPDO@cRGD-M. (C) Drug release profiles of CDDP and OLA from TAT-mPDO@cRGD-M under low-pH and NIR irradiation. (D) Assessment of in vitro hemolysis of TAT-mPDO@cRGD-M (N = 3). (E) In vivo photothermal effect of TAT-mPDO@cRGD-M under NIR irradiation at 2 W/cm^2^ and 1 W/cm^2^. (F) Temperature variation of TAT-mPDO@cRGD-M under NIR irradiation in vitro. (G) SDS-PAGE analysis of membrane proteins derived from MNNG/HOS-CDDP cell membranes (lane 1), cRGD-M (lane 2) and TAT-mPDO@cRGD-M (lane 3).

Dynamic light scattering (DLS) analysis showed that the hydrodynamic diameter of mPDO was approximately 80 nm and increased to ∼100 nm following membrane coating (Fig. 2B). The nanoparticles exhibited a narrow size distribution (Fig. S2B), with polydispersity index (PDI) values below 0.3, indicating good uniformity and reproducibility. No statistically significant difference in zeta potential was observed between TAT-mPDO@cRGD-M and cRGD-M vesicles (Fig. 2B), further confirming successful membrane coating. Drug loading and encapsulation efficiencies of CDDP and OLA in mPDO were quantified by HPLC (Fig. S2A), with loading efficiencies of 17.1% and 22.32%, and encapsulation efficiencies of 20.63% and 28.73%, respectively (Table S4).

Colloidal stability was evaluated in physiological saline and fetal bovine serum (FBS) to simulate isotonic and systemic circulation conditions. No significant changes in particle size or absorbance (Fig. S2G) were observed over 48 h, indicating good structural stability. Hemolysis assays demonstrated negligible hemolytic activity across a range of concentrations (Fig. 2D). In addition, live/dead staining confirmed that neither NIR irradiation alone nor the nanocarrier without drug loading induced measurable cytotoxicity (Fig. S2C), indicating favorable biosafety.

TAT-mPDO@cRGD-M exhibited pronounced NIR responsiveness and pH-dependent drug release behavior. Both acidic conditions and NIR irradiation significantly accelerated the release of CDDP and OLA (Fig. 2C). These effects are consistent with NIR-induced localized thermal effects that may increase membrane permeability and enhance diffusion kinetics of drug molecules within the mesoporous mPDA matrix, thereby facilitating drug release. Additionally, structural modulation of mPDA under acidic conditions may further contribute to accelerated release. The photothermal performance remained stable during repeated irradiation cycles in vitro (Fig. 2F). In vivo evaluation showed that 24 h after intravenous administration, NIR irradiation at 2 W/cm² for 2 min elevated the tumor temperature to above 50 °C (Fig. 2E), confirming efficient photothermal conversion capability.

### Targeting and In Vitro Anti-Tumor Activity of TAT-mPDO@cRGD-M

In the mPDO and TAT-mPDO groups, fluorescence signals were broadly distributed throughout the abdominal region of the nude mice (Fig. 3A), indicating that TAT, as a cell-penetrating peptide, alone is insufficient to achieve in vivo tumor targeting. In contrast, both mPDO@cRGD-M and TAT-mPDO@cRGD-M groups showed significantly improved tumor targeting (Fig. 3C). Ex vivo fluorescence imaging of excised tumors and major organs from nude mice clearly revealed the in vivo biodistribution of TAT-mPDO@cRGD-M (Fig. S2E, F), demonstrating that this nanodelivery system effectively reduced drug accumulation in vital organs, thereby minimizing systemic toxicity. Internalization of drug molecules into the cytoplasm does not necessarily lead to interaction with their subcellular targets^32^—for example, only about 1-4% of cisplatin that is endocytosed enters the nucleus^33^. Confocal microscopy revealed that, following TAT modification, mPDO (green fluorescence, FITC) showed significantly enhanced nuclear localization (blue fluorescence, DAPI) (Fig. 3B, D). Surface modification of the nanoparticle core with TAT peptides thus enabled nuclear-targeted delivery of the nanodrug, bridging the “last mile” from the entry of CDDP into the body to its engagement with tumor DNA. It should be noted that cargos transported via facilitated diffusion through the nuclear pore complex generally have a maximum diameter of approximately 40 nm. In contrast to classical nuclear localization signals (cNLSs), TAT peptides have been reported to promote the nuclear entry of larger and structurally rigid nanoparticles (>40 nm)^34^. The precise mechanism underlying this phenomenon has not yet been fully elucidated.

All in vitro functional assays were performed under the premise of NIR irradiation administered after 4 hours of nanodrug treatment. In the live/dead cell staining assay, the decreased cell viability in the mPD group was attributed to photothermal therapy, while cell viability in the mPDO group was significantly reduced compared to the mPD group (Fig. 3E, H). CDDP-induced DNA damage results in single-strand breaks (SSBs), while PARP1 acts as a key enzyme in their repair. CDDP-resistant cells possess a higher capacity for DNA damage repair compared to wild-type cells; therefore, inhibition of PARP1 using olaparib leads to the accumulation of SSBs, which are subsequently converted into double-strand breaks (DSBs)^35^, thereby reversing CDDP resistance. Under identical drug concentration and treatment duration, the TAT-mPDO@cRGD-M group exhibited the strongest antitumor effect, and both increasing drug concentration and prolonging exposure time further enhanced the tumoricidal activity of the nanodrug (Fig. 3G). Surface modification with either TAT or cRGD further augmented the cytotoxicity of the nanodelivery system toward cancer cells, possibly due to TAT-mediated tumor penetration^36^ or cRGD-facilitated cellular uptake via integrin receptor-mediated endocytosis^37^, which increased intracellular drug accumulation efficiency. TAT-mPDO@cRGD-M exhibited the strongest inhibition of cell migration (Fig. 3E, F). The above experiments were repeated using another CDDP-resistant cell line, U2OS-CDDP, and yielded consistent results (Fig. S3).

**Fig. 3.**
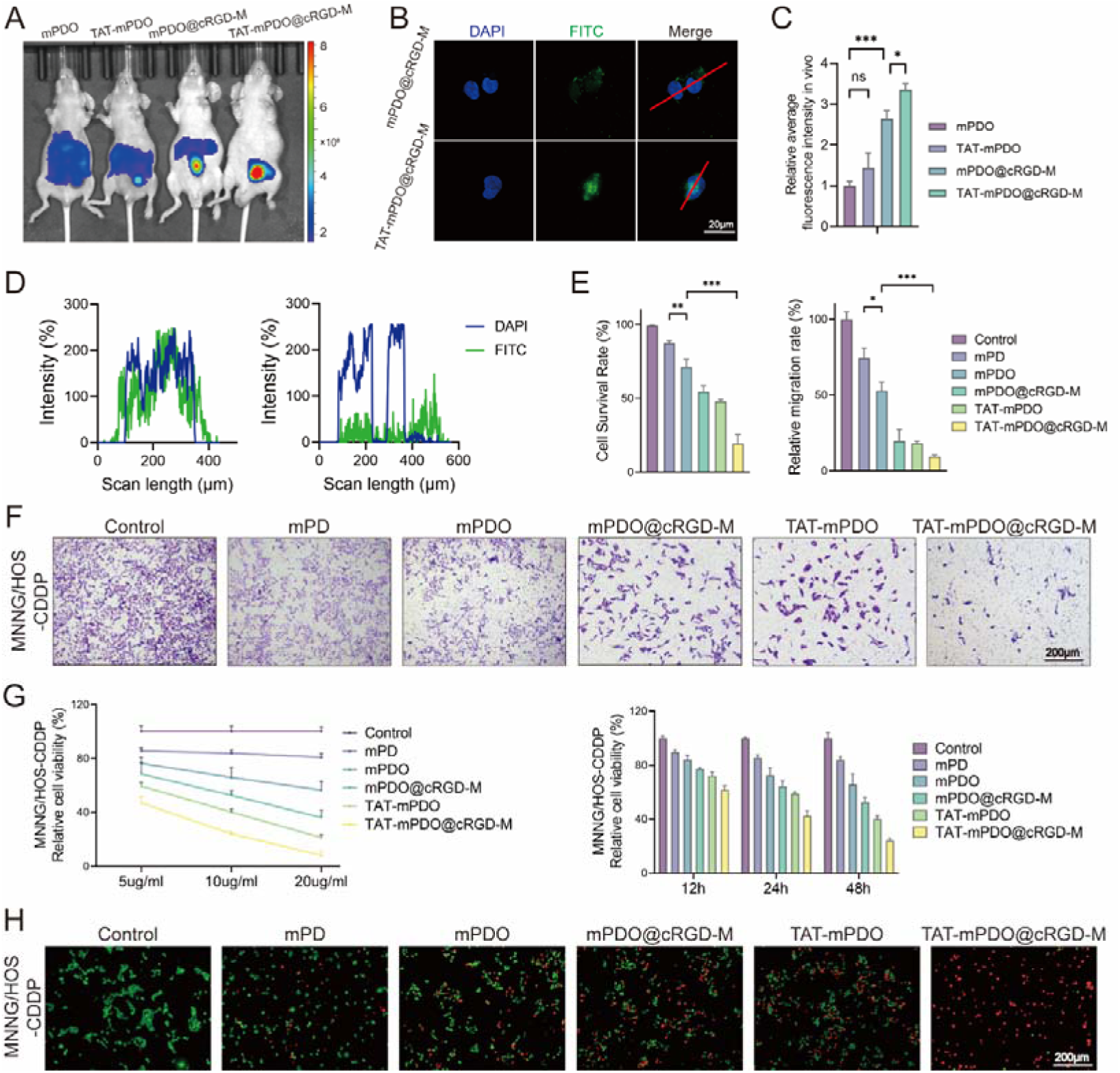
Tumor-targeting capability and antitumor efficacy in vitro of TAT-mPDO@cRGD-M. (A) Biodistribution of mPDO, TAT-mPDO, mPDO@cRGD-M, and TAT-mPDO@cRGD-M in subcutaneous tumor-bearing mice and (C) quantitative analysis (ns: no significance, *p < 0.05, ***p < 0.001, N = 3). (B) mPDO was labeled with FITC (green) and the cell nucleus was labeled with DAPI (blue). The tumor cell nucleus-targeting ability of TAT-mPDO@cRGD-M was investigated in vitro (scale bar: 20 μm). (D) Colocalization analysis of FITC and DAPI fluorescence signals in different groups in vitro. (E) Statistical analysis of live/dead staining and transwell assays (n = 3, mean ± SD, *p < 0.05, **p < 0.01, ***p < 0.001). (F) Transwell assay demonstrating the inhibitory effect of TAT-mPDO@cRGD-M on the migration of MNNG/HOS-CDDP cells (scale bar: 200 μm). (G) CCK-8 assay validating the cytotoxic effects of TAT-mPDO@cRGD-M on MNNG/HOS-CDDP cells at different concentrations and incubation times. (H) Live/dead staining assay validating the cytotoxic effect of TAT-mPDO@cRGD-M on MNNG/HOS-CDDP cells (scale bar: 200 μm).

### Mechanisms of CDDP Resistance Reversal by TAT-mPDO@cRGD-M

To explore the molecular basis underlying the anti-tumor mechanism of TAT-mPDO@cRGD-M, transcriptomic sequencing was performed. A total of 533 DEGs were identified between the TAT-mPDO@cRGD-M and control groups, including 162 upregulated and 371 downregulated genes (Fig. 4A, B). GO enrichment analysis revealed significant clustering of DEGs in biological processes associated with regulation of angiogenesis and regulation of epithelial cell migration, while KEGG pathway analysis identified the PI3K-Akt signaling pathway as one of the most enriched (Fig. 4D). KEGG analysis with LogFC showed an overall suppression trend within these pathways (Fig. 4E). GSEA further demonstrated significant enrichment of the apoptosis hallmark gene set, supporting activation of apoptotic programs upon treatment (Fig. 4C).

**Fig. 4.**
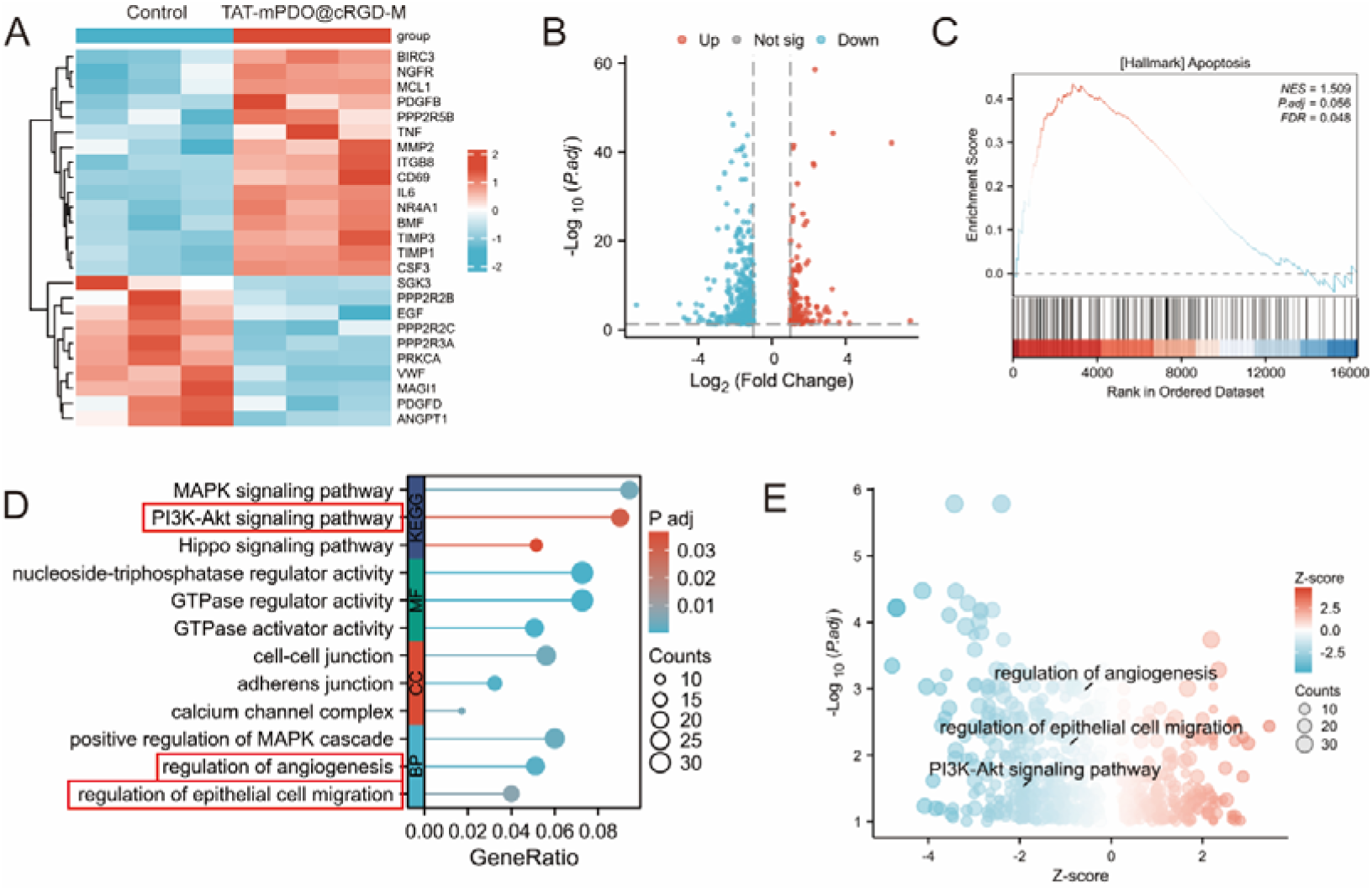
Transcriptomic profiling reveals pathway modulation induced by TAT-mPDO@cRGD-M. (A) Heatmap of PI3K-AKT signaling, apoptosis, and EMT-related genes between the control and the TAT-mPDO@cRGD-M group. (B) Volcano plot of DEGs between the control and the TAT-mPDO@cRGD-M group. (C) GSEA identifying enrichment of apoptosis. (D) GO and KEGG enrichment analysis of DEGs in the TAT-mPDO@cRGD-M group. (E) Integrated GO/KEGG enrichment and LogFC analyses showing three downregulated pathways.

Based on the transcriptomic evidence of apoptosis activation, intracellular ROS levels were first evaluated. Flow cytometric analysis using DCF-FITC staining revealed a concentration-dependent increase in ROS accumulation following TAT-mPDO@cRGD-M treatment (Fig. S4A), suggesting enhanced oxidative stress. Consistently, Annexin V-FITC/PI staining demonstrated a marked elevation in apoptotic cell populations. In MNNG/HOS-CDDP cells, the proportion of early apoptotic cells was substantially increased in the mPDO-treated group relative to control and mPD groups (Fig. S4B), indicating effective induction of programmed cell death.

At the transcriptional level, qPCR analysis revealed significant downregulation of key PI3K-Akt pathway components, including PIK3C3, AKT isoforms, MTOR, and RPTOR, following TAT-mPDO@cRGD-M treatment (Fig. S4C). Consistently, Western blot analysis demonstrated reduced phosphorylation of PI3K and AKT, as reflected by decreased p-PI3K/PI3K and p-AKT/AKT ratios (Fig. 5E, H). Concomitant with PI3K-Akt suppression, apoptosis-related proteins were markedly altered. p-H2AX, BAX, Cleaved Caspase-3, and Cleaved PARP1 were significantly elevated, whereas BCL-2 protein levels were reduced (Fig. 5B, F). qPCR results further confirmed BAX upregulation without significant changes in BCL-2 transcripts (Fig. S4C). Given the enrichment of epithelial migration-related processes, EMT-associated markers were subsequently assessed. FN1, VIM, CDH2, and MMP9 mRNA levels were decreased, accompanied by upregulation of CDH1 (Fig. S4C). Protein expression of N-cadherin, vimentin, and MMP3 was likewise reduced (Fig. 5C, G). To further evaluate the functional involvement of PI3K-Akt signaling, cells were pretreated with 20 μM 740Y-P for 1 h prior to nanodrug administration. 740Y-P partially restored the reduced p-PI3K and p-AKT levels induced by TAT-mPDO@cRGD-M. This restoration reduced p-H2AX accumulation, decreased activation of apoptosis-related proteins, and partially reversed EMT-associated protein changes (Fig. 5D, J). Quantitative analysis was consistent with immunoblot observations.

**Fig. 5.**
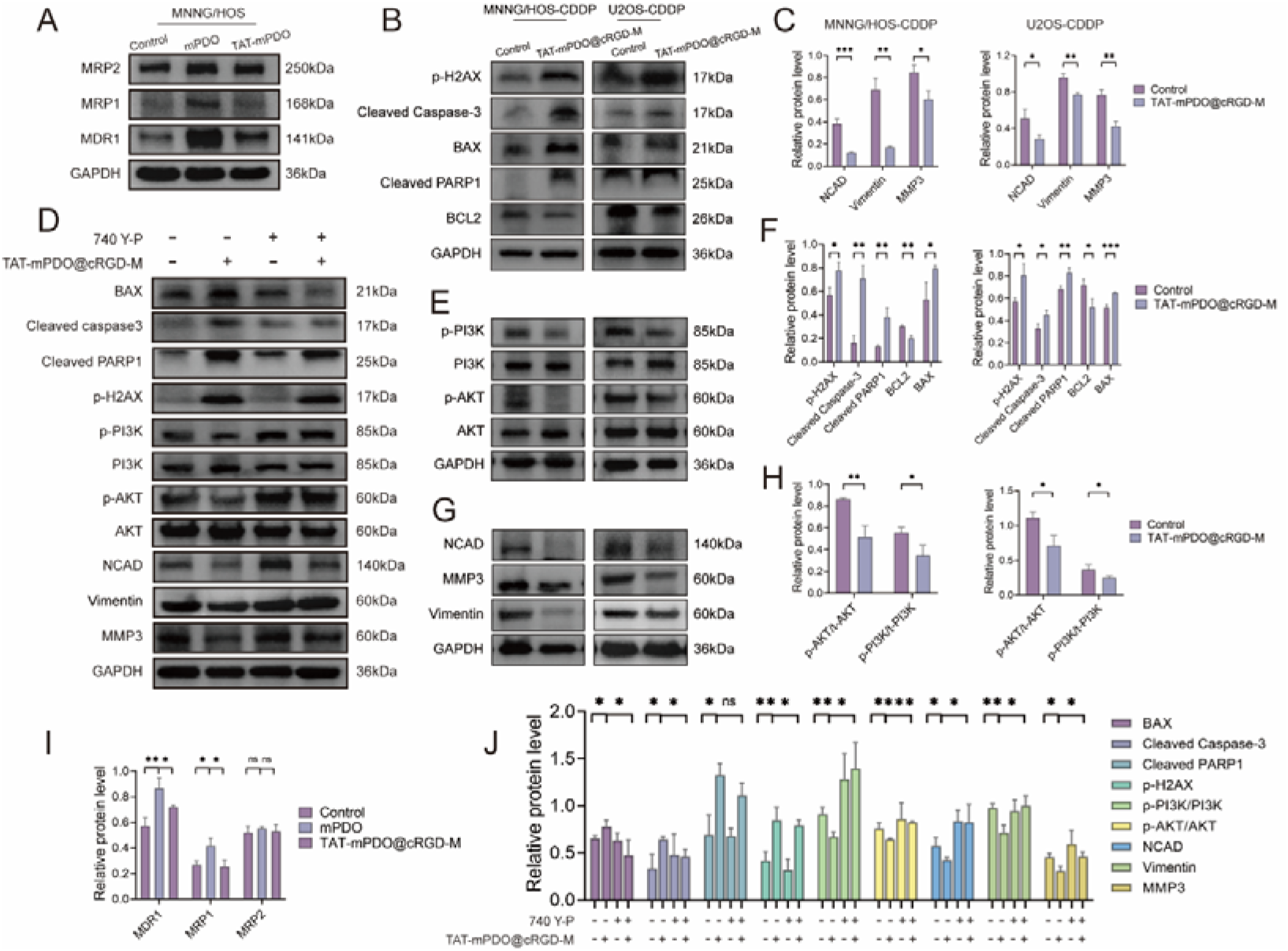
TAT-mPDO@cRGD-M suppresses PI3K-AKT signaling to promote apoptosis and inhibit EMT in resistant osteosarcoma. (A) Western blot analysis of MRP1/2 and MDR1 expression in MNNG/HOS cells from the mPDO and TAT-mPDO groups. (B) Western blot analysis of apoptosis-related proteins, key proteins in the PI3K–AKT signaling pathway (E), and EMT-associated proteins (G) in MNNG/HOS-CDDP and U2OS-CDDP cells after treatment with TAT-mPDO@cRGD-M. (C) Quantitative analysis of the proteins in (G) (n = 3, mean ± SD, *p < 0.05, **p < 0.01, ***p < 0.001). (D) Western blot analysis in MNNG/HOS-CDDP cells assessing the effects of 740Y-P and TAT-mPDO@cRGD-M treatment on apoptosis- and PI3K-AKT/EMT-associated proteins. (F) Quantitative analysis of the proteins in (B) (n = 3, mean ± SD, *p < 0.05, **p < 0.01, ***p < 0.001). (H) Quantitative analysis of the proteins in (E) (n = 3, mean ± SD, *p < 0.05, **p < 0.01). (I) Quantitative analysis of the proteins in (A) (n = 3, mean ± SD, ns: no significance, *p < 0.05, **p < 0.01). (J) Summary quantification of representative proteins indicated in (D) (n = 3, mean ± SD, ns: no significance, *p < 0.05, **p < 0.01, ***p < 0.001).

**Fig. 6.**
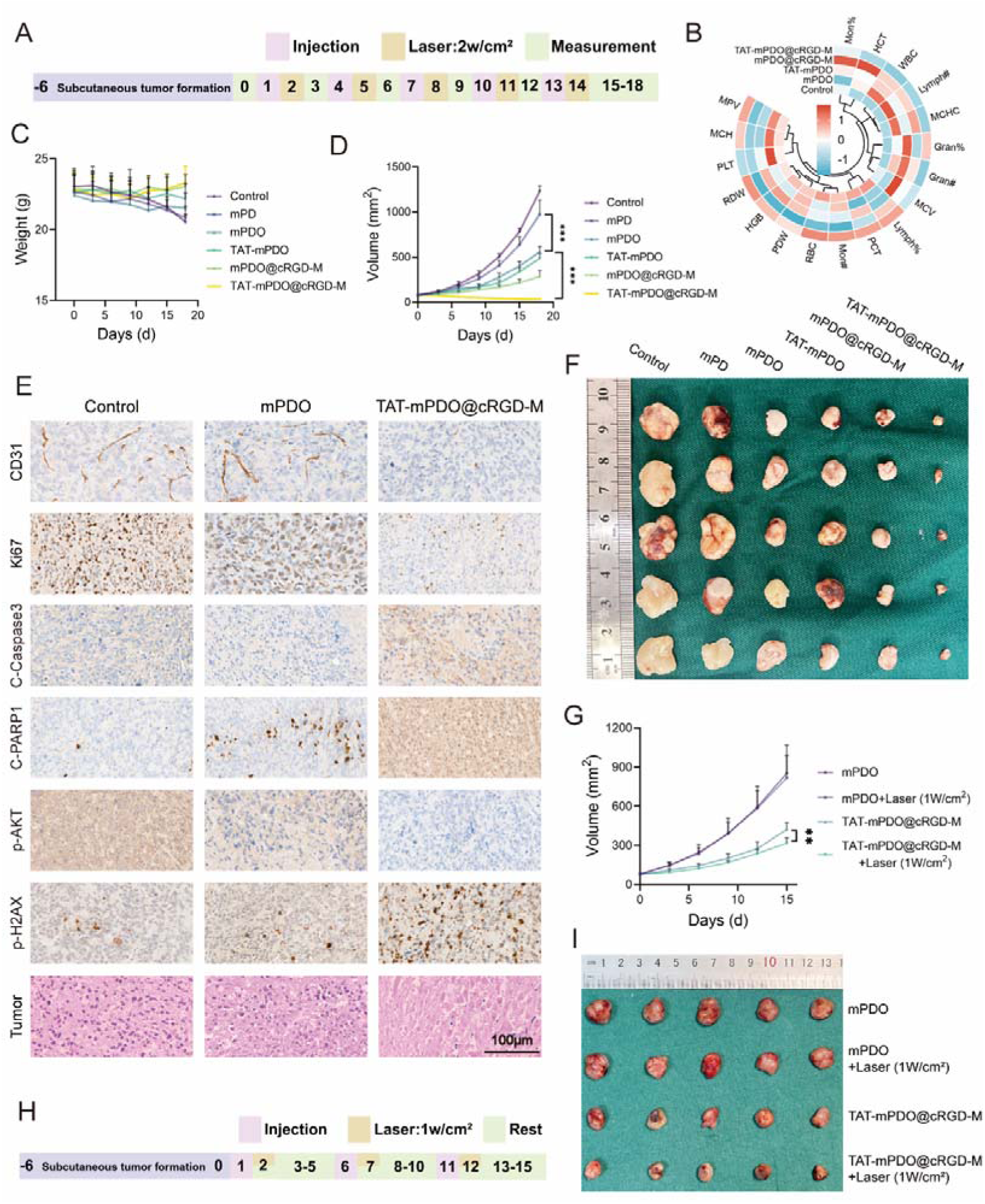
Anti-tumor efficacy of TAT-mPDO@cRGD-M in vivo. (A) Experimental timeline schematic of the in vivo experiment using a subcutaneous osteosarcoma xenograft model. (B) Heatmap of biochemical parameters in peripheral blood from nude mice of each group. (C) Nude mice were divided into groups: control, mPD, mPDO, TAT-mPDO, mPDO@cRGD-M, and TAT-mPDO@cRGD-M. Body weight of nude mice was recorded over time. (D) Tumor growth curves of above 6 groups (***p < 0.001, N = 5). (E) H&E staining of tumors and IHC staining of CD31, Ki67, C-Caspase3, C-PARP1, p-AKT and p-H2AX in control, mPDO and TAT-mPDO@cRGD-M groups (scale bar: 100 μm). (F) Representative photograph of excised tumors from above 6 groups. (G) Tumor growth curves of groups in (I) (**p < 0.01, N = 5). (H) Schematic illustration of the in vivo experimental procedure under low-power NIR irradiation. (I) Nude mice were divided into groups: mPDO, mPDO+Laser (1 W/cm²), TAT-mPDO@cRGD-M, TAT-mPDO@cRGD-M+Laser (1 W/cm²) and the representative photograph of excised tumors from these groups

Drug efflux mediated by ABC transporters is widely implicated in acquired chemoresistance^38,39^. We therefore evaluated the expression of major multidrug resistance proteins, including MRP1, MRP2, and MDR1. In wild-type MNNG/HOS cells, mPDO treatment markedly increased the protein levels of MRP1 and MDR1, whereas MRP2 expression showed no significant change. Notably, TAT-mPDO treatment significantly attenuated the upregulation of MRP1 and MDR1 observed in the mPDO group, while MRP2 remained unaffected (Fig. 5A, I). These findings suggest that TAT modification selectively attenuates the adaptive upregulation of specific efflux transporters, which enhances intracellular drug retention^34^.

### In Vivo Tumor Suppression and Mild Photothermal Therapy

A subcutaneous xenograft nude mouse model was established using the MNNG/HOS-CDDP cell line to investigate the in vivo antitumor efficacy of the nanodrug delivery system against CDDP-resistant osteosarcoma (Fig. 6A). In the mPDO group, tumor volume was markedly reduced (Fig. 6D), indicating that the strategy of reversing CDDP resistance with olaparib is also effective in vivo. TAT-mPDO did not exhibit enhanced antitumor activity compared with mPDO. In contrast, TAT-mPDO@cRGD-M showed significantly greater tumor growth inhibition than mPDO@cRGD-M (Fig. 6F, S5B), which may be because TAT alone has limited tumor-targeting ability in vivo and depends on cRGD to direct it into tumor cells before acting on the nucleus. Hematoxylin-eosin (H&E) staining revealed that tumors treated with the nanodrug exhibited disorganized cellular arrangement, cytoplasmic vacuolation, and necrosis or fibrosis (Fig. 6E). No obvious signs of cell necrosis, foamy degeneration, or coagulative necrosis were observed in vital organs, and tissue structures remained intact (Fig. S5A). Body weight changes during treatment were minimal in the TAT-mPDO@cRGD-M group (Fig. 6C), peripheral blood RBC, WBC, and platelet counts and morphology did not differ significantly from the control group (Fig. 6B). Overall, these results demonstrate that the TAT-mPDO@cRGD-M nano-drug delivery system produces significant suppression of subcutaneous tumor growth while exhibiting high biosafety. Immunohistochemical analysis further showed increased staining for Cleaved Caspase-3, Cleaved PARP1, and p-H2AX, along with decreased Ki-67 and p-AKT in the TAT-mPDO@cRGD-M group (Fig. 6E), supporting that this nano-delivery system suppresses the PI3K-AKT signaling pathway and promotes tumor cell apoptosis. Since α_v_β_3_ is highly expressed in tumor neovasculature^28^, immunohistochemistry revealed reduced staining of the neovascular marker CD31 (Fig. 6E), suggesting that cRGD-functionalized nanoparticles not only target tumor cells but also damage endothelial cells in newly formed vessels, thereby inhibiting tumor growth by disrupting the nutrient supply.

mPD combined with photothermal therapy (PTT) inhibited the growth of cisplatin-resistant tumors to some extent (Fig. 6D), likely due to the ability of PTT to directly induce tumor cell damage through localized hyperthermia, bypassing chemotherapy resistance. Conventional PTT typically elevates the local temperature above 50 °C for effective tumor ablation, but such high temperatures may damage surrounding normal tissues^40^. To minimize the impact on normal tissues, mild PTT maintains the temperature between 42°C and 45°C^41^. Within this range, heat shock protein expression in tumor cells can be effectively suppressed, reducing thermotolerance, thereby enhancing therapeutic outcomes while minimizing injury to normal tissues^42^. Based on this rationale, we investigated the in vivo anticancer efficacy of the nano-delivery system under low-power NIR irradiation (Fig. 6H). At 24 h post-injection, NIR irradiation at 1 W/cm² resulted in a maximum tumor temperature of approximately 45 °C (Fig. 2E), which caused notably less cutaneous burn damage than 2 W/cm² irradiation (Fig. S2D). However, 1 W/cm² NIR combined with mPDO did not show synergistic effects, possibly due to insufficient tumor accumulation of mPDO and limited thermal diffusion, which hindered heat transfer to the deep DNA in the nucleus^34^. In contrast, tumor volume (Fig. 6G, I) and weight (Fig. S5C) were significantly lower in the TAT-mPDO@cRGD-M + Laser (1 W/cm²) group than in the TAT-mPDO@cRGD-M group. These findings suggest that the cascade delivery system can achieve effective photothermal therapy at lower power densities and temperatures by generating localized hyperthermia near nuclear DNA.

### Conclusion

This study developed a biomimetic nanodrug system, TAT-mPDO@cRGD-M, offering an innovative therapeutic strategy for cisplatin-resistant osteosarcoma. TAT-mPDO@cRGD-M enhances DNA damage and suppresses the PI3K-AKT signaling pathway, promoting apoptosis and inhibiting epithelial-mesenchymal transition, thereby effectively preventing tumor cell proliferation and migration. In vivo, TAT-mPDO@cRGD-M significantly inhibited tumor growth and sustained antitumor activity even under mild-temperature conditions, without causing noticeable damage to normal tissues. Overall, TAT-mPDO@cRGD-M shows great promise in overcoming chemoresistance in osteosarcoma and has substantial potential for future clinical applications.

## Authors’ contributions

XYL, SJX and YYP: Conceptualization, Investigation, Methodology, Software, Writing original draft. ZLS, SNW and AL: Supervision, Data curation. HX, ZWS and YYD: Funding acquisition, Supervision, Review & Editing. All authors have given final approval for this version of the manuscript to be published.

## Ethics declarations

This study was approved by the Medical Ethics Committee of Union Hospital of Huazhong University of Science and Technology (Approval number: [2023] Ethics Review (0046)).

All animal experiments were approved by the Institutional Animal Care and Use Committee of Huazhong University of Science and Technology (Approval number: [2024] IACUC number: 4351) and conducted in accordance with institutional and national ethical guidelines.

## Acknowledgments

The authors would like to express their gratitude to all collaborators and colleagues who provided valuable insights and constructive discussions during the preparation of this article. Additionally, we acknowledge the support from Huazhong University of Science and Technology for providing access to relevant literature.

