## supplementary figure for "Biomimetic Cascade-Targeting Drug Delivery System for Reversing Chemoresistance in Osteosarcoma"

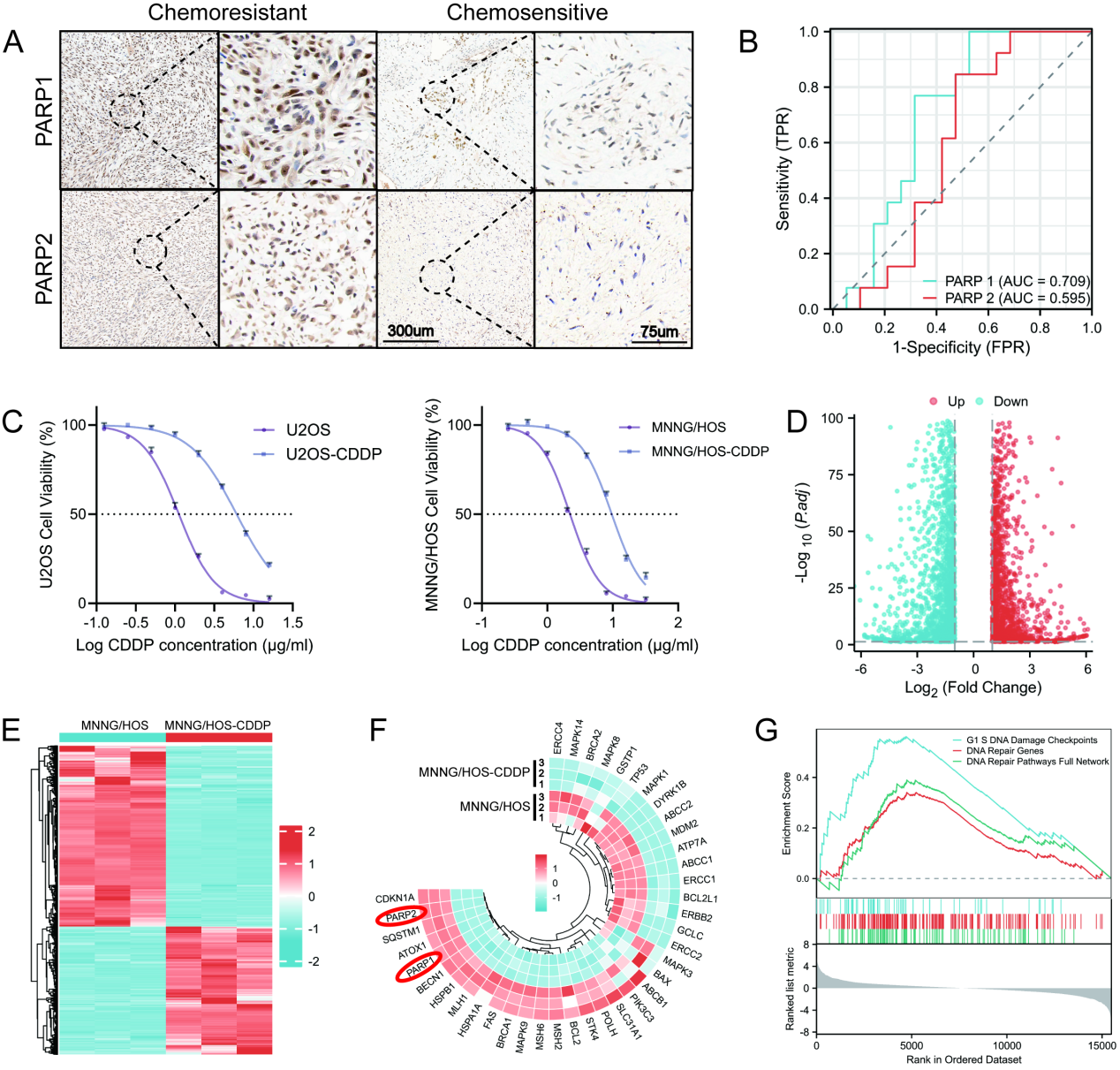

**Fig. S1 PARP1/2 expression and molecular characterization of CDDP-resistant osteosarcoma.**

(A) Immunohistochemical expression of PARP1/2 in chemoresistant and chemosensitive osteosarcomas (scale bar: 300 μm, 75 μm).
    (B) ROC curve of PARP1/2 for predicting chemotherapy sensitivity in osteosarcoma.
    (C) Construction of cisplatin-resistant MNNG/HOS-CDDP and U2OS-CDDP cell lines and measurement of IC_50_ values.
    (D) Volcano plot of DEGs between MNNG/HOS and MNNG/HOS-CDDP cells.
    (E) Heatmap of DEGs between MNNG/HOS and MNNG/HOS-CDDP cells.
    (F) Expression differences of cisplatin resistance-associated genes between MNNG/HOS and MNNG/HOS-CDDP cells.
    (G) GSEA revealing enrichment of DNA damage repair-related pathways.

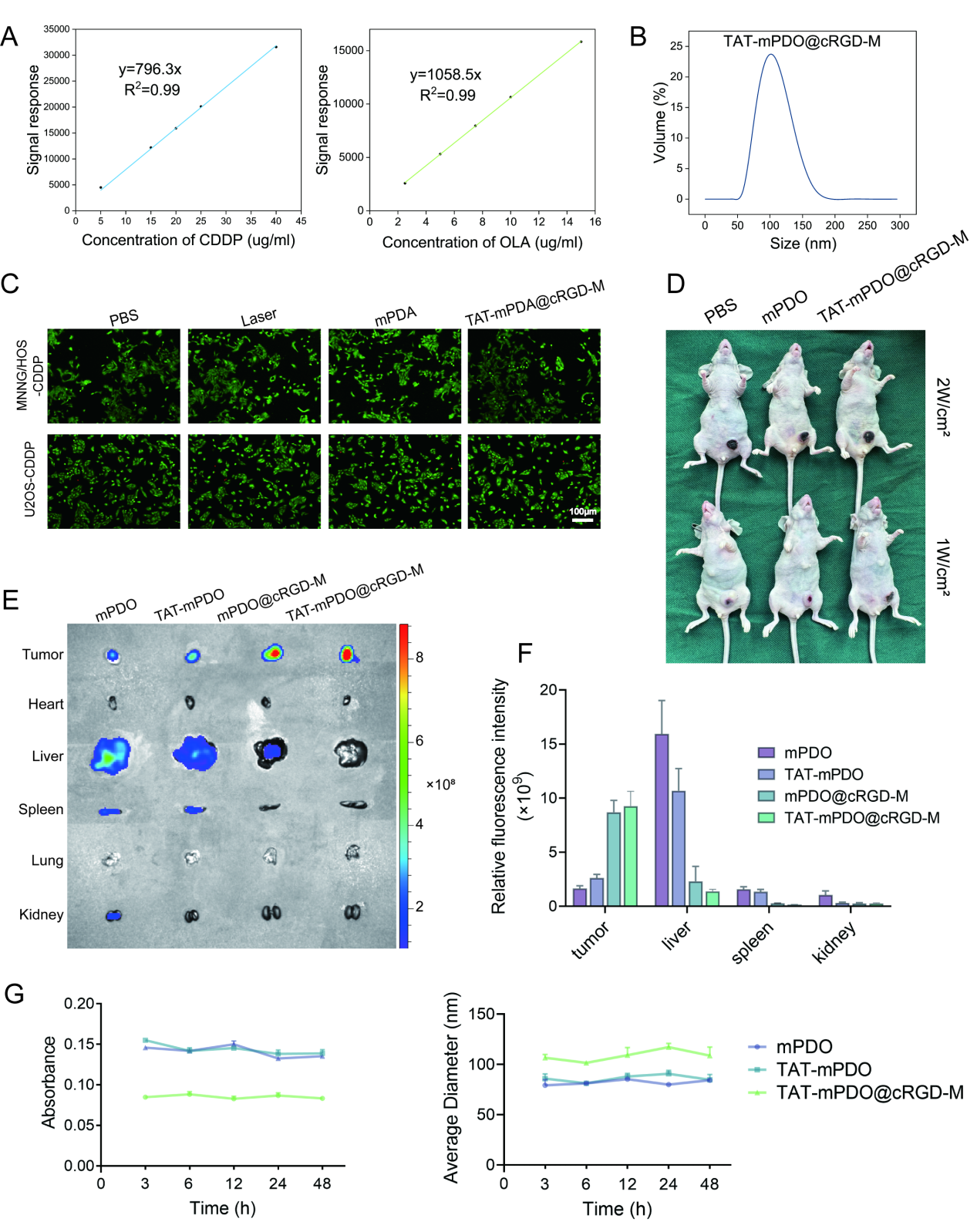

**Fig. S2 Characterization and drug distribution of TAT-mPDO@cRGD-M.**

(A) Standard curves of CDDP and OLA.
(B) Particle size distribution of TAT-mPDO@cRGD-M.
(C) Cytotoxicity assays of mPDA, TAT-mPDA@cRGD-M and NIR irradiation on cells (scale bar: 100 µm).
(D) Impact of the nanodelivery system on peritumoral skin and normal tissues under NIR irradiation at different power.
(E) Uptake of drugs in the heart, liver, spleen, lung, and kidney of different groups, as well as drug distribution in tumors and (F) statistical analysis (N = 3).

(G) The absorbance changes of mPDO, TAT-mPDO, and TAT-mPDO@cRGD-M in FBS over 48h and the particle size variation in physiological saline over 48h.

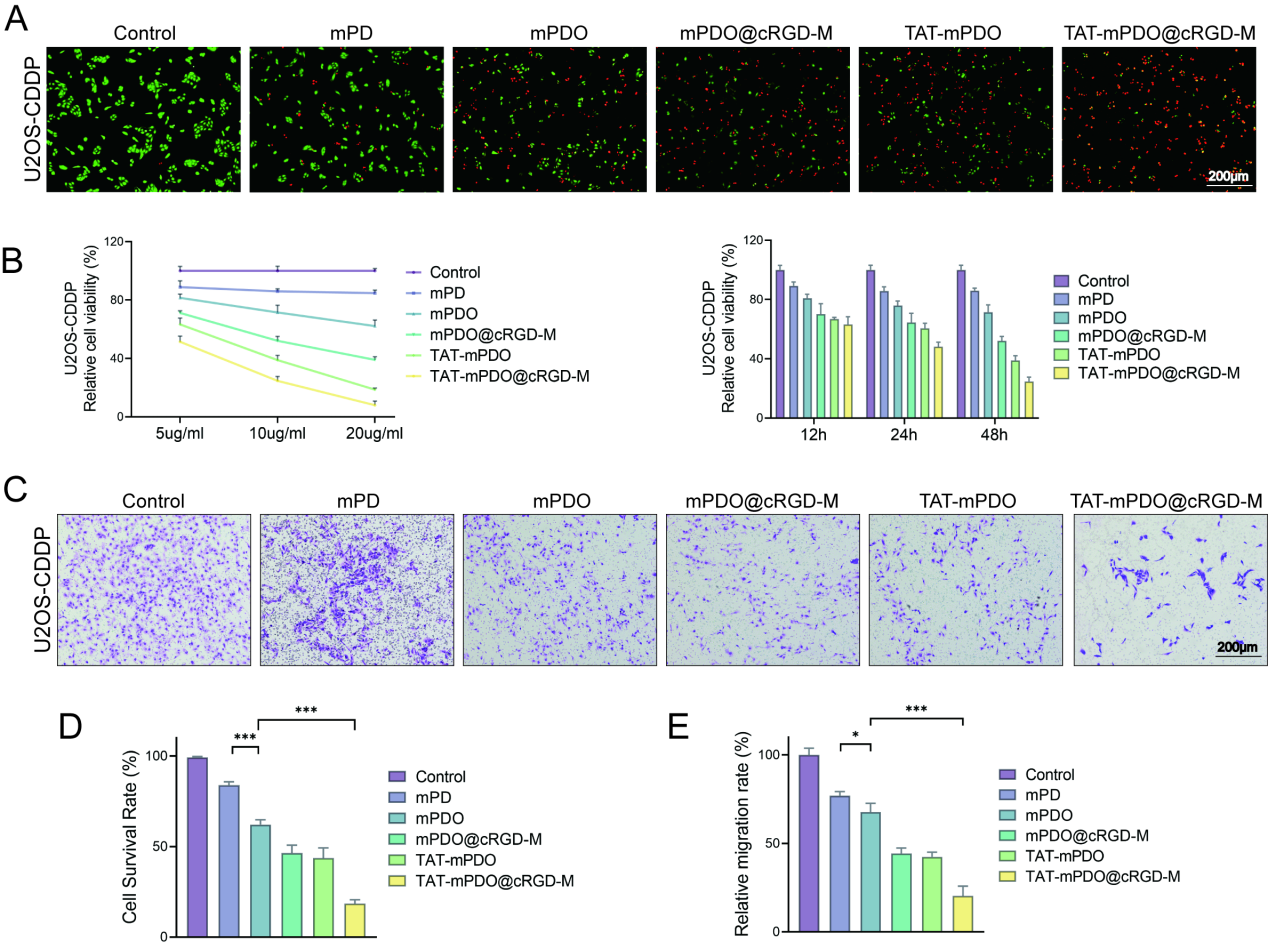

**Fig. S3 Antitumor efficacy of TAT-mPDO@cRGD-M U2OS-CDDP in vitro.**

(A) Live/dead staining assay validating the cytotoxic effect of TAT-mPDO@cRGD-M on U2OS-CDDP cells (scale bar: 200 μm).
    (B) CCK-8 assay validating the cytotoxic effects of TAT-mPDO@cRGD-M on U2OS-CDDP cells at different concentrations and incubation times.
    (C) Transwell assay demonstrating the inhibitory effect of TAT-mPDO@cRGD-M on the migration of U2OS-CDDP cells (scale bar: 200 μm).
    (D, E) Statistical analysis of live/dead staining and transwell assays (n = 3, mean ± SD, *p < 0.05, ***p < 0.001).

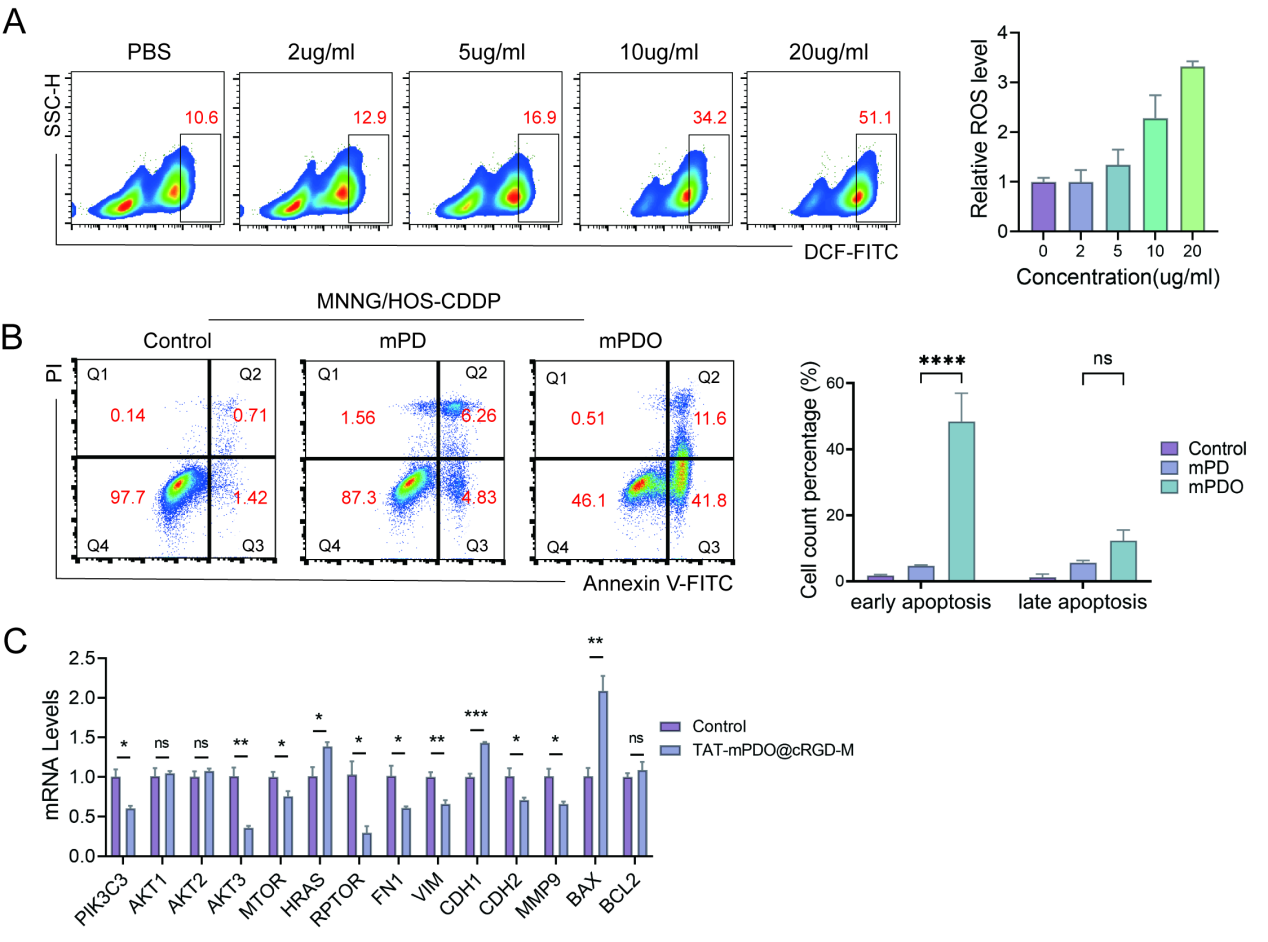

**Fig. S4 Anti-tumor mechanism analysis of TAT-mPDO@cRGD-M treatment.**

(A) Flow cytometry analysis of intracellular ROS levels in cells treated with different concentrations of TAT-mPDO@cRGD-M and statistical analysis (N = 3).
    (B) Flow cytometric analysis of apoptosis in the mPDO group and statistical analysis (ns: no significance, ****p < 0.0001, N = 3).
    (C) RT-qPCR analysis of the expression of genes related to the PI3K-AKT signaling pathway, apoptosis and EMT in the TAT-mPDO@cRGD-M group.

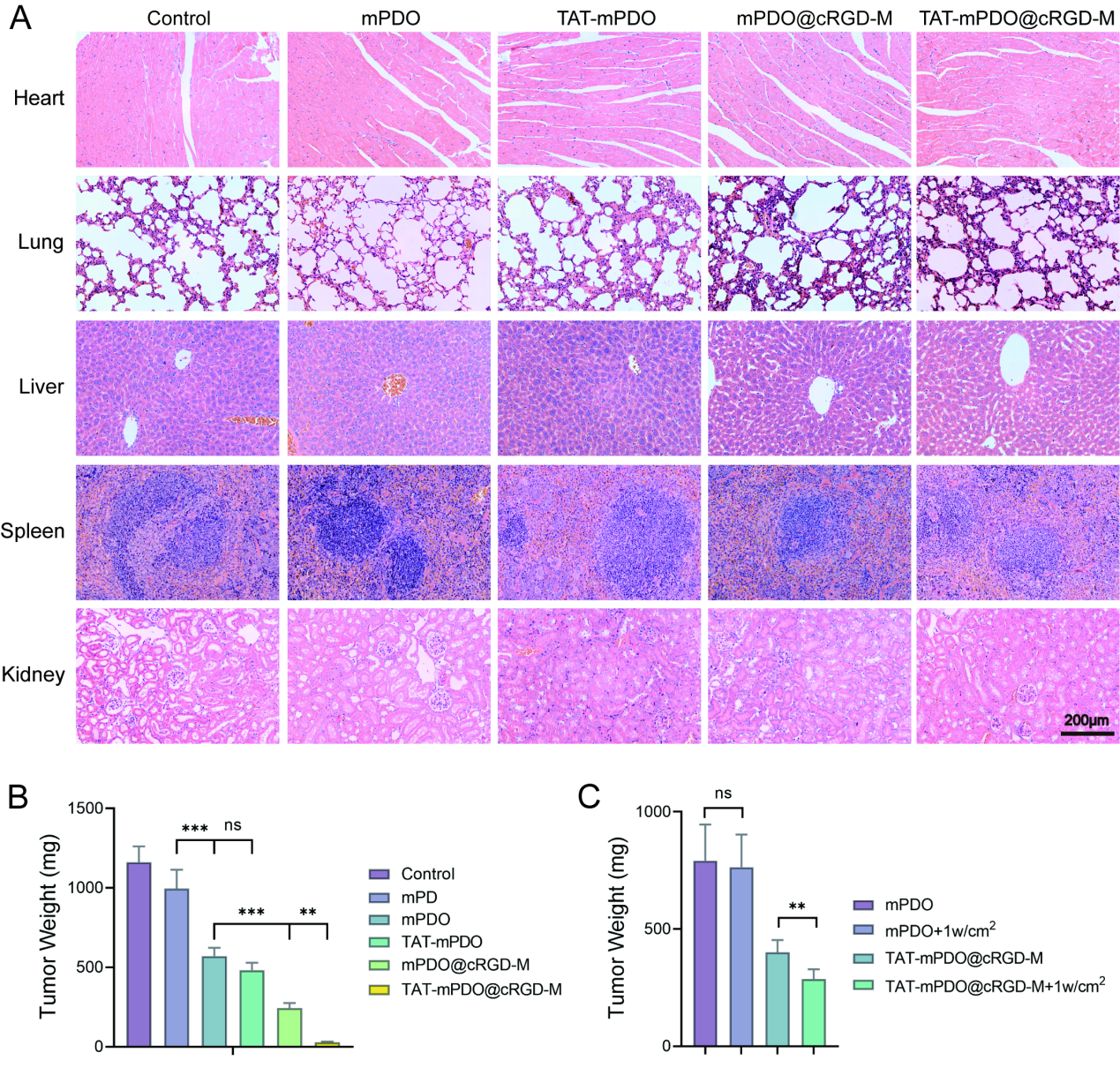

**Fig. S5 In vivo toxicity and anti-tumor efficacy of TAT-mPDO@cRGD-M.**

(A) H&E staining of major organs (heart, lung, liver, spleen and kidney) from nude mice in the control, mPDO, TAT-mPDO, mPDO@cRGD-M, and TAT-mPDO@cRGD-M groups.
(B) Final tumor weight from the control, mPD, mPDO, TAT-mPDO, mPDO@cRGD-M, and TAT-mPDO@cRGD-M groups (ns: no significance, **p < 0.01, ***p < 0.001, N = 5).
(C) Final tumor weight from the mPDO, mPDO+Laser (1 W/cm²), TAT-mPDO@cRGD-M, TAT-mPDO@cRGD-M+Laser (1 W/cm²) groups (ns: no significance, **p < 0.01, N = 5).
