## supplementary table for "Biomimetic Cascade-Targeting Drug Delivery System for Reversing Chemoresistance in Osteosarcoma"

| **Table S1** | Mapping rate of six samples in group MNNG/HOS and MNNG/HOS-CDDP |
| --- | --- |
| **Table S2** | Mapping rate of six samples in group control and TAT-mPDO@cRGD-M |
| **Table S3** | Primer sequences used in this study |
| **Table S4** | The calculation process of the loading efficiency and encapsulation efficiency of nanodrugs |

Table S1.

| sample | total_reads | total_map | unique_map | multi_map | read1_map | read2_map | positive_map | negative_map | splice_map | unsplice_map | proper_map |
| --- | --- | --- | --- | --- | --- | --- | --- | --- | --- | --- | --- |
| MNNG/HOS1 | 42826778 | 40587250(94.77%) | 39511302(92.26%) | 1075948(2.51%) | 19783394(46.19%) | 19727908(46.06%) | 19729533(46.07%) | 19781769(46.19%) | 17910460(41.82%) | 21600842(50.44%) | 37173374(86.8%) |
| MNNG/HOS2 | 43009340 | 41009024(95.35%) | 39950963(92.89%) | 1058061(2.46%) | 20028189(46.57%) | 19922774(46.32%) | 19947399(46.38%) | 20003564(46.51%) | 17725775(41.21%) | 22225188(51.68%) | 37988136(88.33%) |
| MNNG/HOS3 | 42861842 | 40789102(95.16%) | 39725046(92.68%) | 1064056(2.48%) | 19917600(46.47%) | 19807446(46.21%) | 19836094(46.28%) | 19888952(46.4%) | 17858341(41.66%) | 21866705(51.02%) | 37736354(88.04%) |
| MNNG/HOS-CDDP1 | 42705706 | 40797414(95.53%) | 39695125(92.95%) | 1102289(2.58%) | 19894886(46.59%) | 19800239(46.36%) | 19820269(46.41%) | 19874856(46.54%) | 18637980(43.64%) | 21057145(49.31%) | 37912430(88.78%) |
| MNNG/HOS-CDDP2 | 44093144 | 41839587(94.89%) | 40658267(92.21%) | 1181320(2.68%) | 20378864(46.22%) | 20279403(45.99%) | 20296994(46.03%) | 20361273(46.18%) | 19520209(44.27%) | 21138058(47.94%) | 38688400(87.74%) |
| MNNG/HOS-CDDP3 | 39955572 | 38214633(95.64%) | 37190978(93.08%) | 1023655(2.56%) | 18638478(46.65%) | 18552500(46.43%) | 18571412(46.48%) | 18619566(46.6%) | 17552925(43.93%) | 19638053(49.15%) | 35453786(88.73%) |

Table S2.

| SampleID | Total reads | Total mapped | Multiple mapped | Uniquely mapped |
| --- | --- | --- | --- | --- |
| Control1 | 55186008 | 53870957 (97.62%) | 1459419 (2.64%) | 52411538 (94.97%) |
| Control2 | 49647672 | 48663142 (98.02%) | 1299620 (2.62%) | 47363522  (95.4%) |
| Control3 | 36693370 | 35901399 (97.84%) | 1067663 (2.91%) | 34833736 (94.93%) |
| TAT-mPDO@cRGD-M1 | 59153944 | 57805008 (97.72%) | 1828692 (3.09%) | 55976316 (94.63%) |
| TAT-mPDO@cRGD-M2 | 57724886 | 56377084 (97.67%) | 1772726 (3.07%) | 54604358 (94.59%) |
| TAT-mPDO@cRGD-M3 | 37125410 | 36236255 (97.6%) | 1127344 (3.04%) | 35108911 (94.57%) |

Table S3.

|  | Primer name | Primer sequence |
| --- | --- | --- |
| 1 | GAPDH-F | TCGGAGTCAACGGATTTGGT |
|  | GAPDH-R | TTCCCGTTCTCAGCCTTGAC |
| 2 | PIK3C3-F | CCTGGAAGACCCAATGTTGAAG |
|  | PIK3C3-R | CGGGACCATACACATCCCAT |
| 3 | AKT1-R | GCCATCATTCTTGAGGAGGAAGT |
|  | AKT1-F | AGCGACGTGGCTATTGTGAAG |
| 4 | AKT2-F | AGGCACGGGCTAAAGTGAC |
|  | AKT2-R | CTGTGTGAGCGACTTCATCCT |
| 5 | AKT3-F | AATGGACAGAAGCTATCCAGGC |
|  | AKT3-R | TGATGGGTTGTAGAGGCATCC |
| 6 | MTOR-F | GCAGATTTGCCAACTATCTTCGG |
|  | MTOR-R | CAGCGGTAAAAGTGTCCCCTG |
| 7 | HRAS-F | TGCTTCAGTTTGAACTACCCTG |
|  | HRAS-R | GCCCAGTGCTGATAGCCAG |
| 8 | RPTOR-F | AATGCTGCAATCGCCTCTTCT |
|  | RPTOR-R | GCCAAAGGTAGGTTCCAGTCTG |
| 9 | FN1-F | GAGAATAAGCTGTACCATCGCAA |
|  | FN1-R | CGACCACATAGGAAGTCCCAG |
| 10 | VIM-F | AGTCCACTGAGTACCGGAGAC |
|  | VIM-R | CATTTCACGCATCTGGCGTTC |
| 11 | CDH1-F | ATTTTTCCCTCGACACCCGAT |
|  | CDH1-R | TCCCAGGCGTAGACCAAGA |
| 12 | CDH2-F | AGCCAACCTTAACTGAGGAGT |
|  | CDH2-R | GGCAAGTTGATTGGAGGGATG |
| 13 | MMP9-R | TCGTCATCGTCGAAATGGGC |
|  | MMP9-F | GGGACGCAGACATCGTCATC |
| 14 | BAX-F | CCAGCCCATGATGGTTCTGAT |
|  | BAX-R | CCCGAGAGGTCTTTTTCCGAG |
| 15 | BCL2-F | CGGTTCAGGTACTCAGTCATCC |
|  | BCL2-R | GGTGGGGTCATGTGTGTGG |

Table S4.

| OLA | Peak Area | Amount (ug/ml) |
| --- | --- | --- |
|  | 0 | 0 |
|  | 2580.315 | 2.5 |
|  | 5310.07 | 5 |
|  | 7957.842 | 7.5 |
|  | 10641.68 | 10 |
|  | 15834.596 | 15 |

| CDDP | Peak Area | Amount (ug/ml) |
| --- | --- | --- |
|  | 0 | 0 |
|  | 4501.073 | 5 |
|  | 12189.36 | 15 |
|  | 15912.743 | 20 |
|  | 20098.376 | 25 |
|  | 31585.934 | 40 |

|  |  | Liquid Chromatography Peak Area | Concentration Derived from Standard Curve | Actual Concentration Before Dilution (100times) | Amount of Drug in Supernatant After Multiplying by Dispersion Volume  (6ml) | Dosing Amount (mg) | Drug Loading Amount | Encapsulation Efficiency |
| --- | --- | --- | --- | --- | --- | --- | --- | --- |
| mPDO | OLA | 3396.987 | 0.003064088 | 0.153204415 | 2.298066225 | 8 | 22.32% | 28.73% |
|  | CDDP | 5940.156 | 0.007333803 | 0.110007042 | 1.65010563 | 8 | 17.10% | 20.63% |
